# Scalable, optically-responsive human neuromuscular junction model reveals convergent mechanisms of synaptic dysfunction in familial ALS

**DOI:** 10.1101/2024.01.11.575304

**Authors:** Daniel Chen, Polyxeni Philippidou, Bianca de Freitas Brenha, Ashleigh E. Schaffer, Helen C. Miranda

## Abstract

Neuromuscular junctions (NMJs) are specialized synapses that mediate communication between motor neurons and skeletal muscles and are essential for movement. The degeneration of this system can lead to symptoms observed in neuromuscular and motor neuron diseases. Studying these synapses and their degeneration has proven challenging. Prior NMJ studies heavily relied upon the use of mouse, chick, or isolated primary human cells, which have demonstrated limited fidelity for disease modeling. To enable the study of NMJ dysfunction and model genetic diseases, we, and others, have developed methods to generate human NMJs from pluripotent stem cells (PSCs), embryonic stem cells, and induced pluripotent stem cells. However, published studies have highlighted technical limitations associated with these complex *in vitro* NMJ models. In this study, we developed a robust PSC-derived motor neuron and skeletal muscle co-culture method, and demonstrated its sensitivity in modeling motor neuron disease. Our method spontaneously and reproducibly forms human NMJs. We developed multiwell-multielectrode array (MEA) parameters to quantify the activity of PSC-derived skeletal muscles, as well as measured the electrophysiological activity of functional human PSC-derived NMJs. We further leveraged our method to morphologically and functionally assess NMJs from the familial amyotrophic lateral sclerosis (fALS) PSCs, *C9orf72* hexanucleotide (G4C2)n repeat expansion (HRE), *SOD1*^A5V^, and *TDP43*^G298S^ to define the reproducibility and sensitivity of our system. We observed a significant decrease in the numbers and activity of PSC-derived NMJs developed from the different ALS lines compared to their respective controls. Furthermore, we evaluated a therapeutic candidate undergoing clinical trials and observed a variant-dependent rescue of functionality of NMJs. Our newly developed method provides a platform for the systematic investigation of genetic causes of NMJ neurodegeneration and highlights the need for therapeutic avenues to consider patient genotype.

## INTRODUCTION

Neuromuscular junctions (NMJs) are highly specialized synapses formed between motor neurons and skeletal muscle fibers to convert electrical stimulation into muscle contraction ^1,2^. The release of acetylcholine from motor neurons into the synaptic cleft initiates action potentials across the muscle’s surface, which causes muscles to contract. Dysfunction, disassembly, or degeneration of NMJs is a pathological characteristic of many neurodegenerative and neuromuscular disorders (NMDs) ^3^. The pathological involvement of neurons and muscles varies amongst NMDs. For example, Amyotrophic Lateral Sclerosis (ALS) and Spinal Muscular Atrophy (SMA) have distinguished neuronal pathology ^3,4^, whereas Muscular Dystrophy (DM) is a known myopathy ^5^, and patients with Spinal Bulbar Muscular Atrophy (SBMA) can have both signs of neurogenic atrophy and myopathy ^6^. Additionally, one key hallmark of aging is the deleterious accumulation of mislocalized acetylcholine receptors (AChR) at the NMJ, which eventually causes its degeneration ^1,2^. Animal models have been developed to investigate the progression of various neuromuscular diseases ^7^. While they have provided valuable information, the caveat to these studies is that the observed events may not accurately mimic what occurs in humans. Indeed, comparative studies between mice and humans have revealed important differences in NMJs, for instance, human NMJs are much smaller and more fragmented than those of mice ^3,8,9^. These species-specific differences could contribute to observed dissimilarities in NMD presentation and disease progression in mice and humans with identical genetic mutations. Moreover, in December of 2022, the FDA Modernization Act 2.0 was signed into law stating that animal testing is no longer mandated for every new drug development ^10^. This legislation underscores the importance of human cell-based models, not only critical to understanding NMJ dysfunction in NMDs, but as a platform for new therapeutic discoveries ^11^.

Patient-specific induced PSCs (iPSCs) allow for the genomes of human subjects afflicted with NMDs to be captured in a pluripotent stem cell line ^12,13^. Interestingly, the first report of iPSC applied to disease modeling was in the context of ALS in 2008 ^14^. Since then, significant advancements have been made in understanding ALS pathology in individual cell types such as motor neurons, astrocytes, microglia, and skeletal muscles ^14–17^. ALS is now considered a “multisystemic” disease, and NMJ degeneration is thought to precede neuronal cell death ^18^. Moreover, therapeutic targets in iPSC-derived NMD models have largely focused on addressing motor neuron phenotypes that would prolong neuronal survival *in vitro* ^4,19,20^. Although initially thought to be promising, the field now acknowledges that preventing motor neuron loss does not assure muscle innervation is preserved, nor does it ensure re-innervation. In recent years, several labs have published NMJ co-culture systems, however, those methods have heavily relied upon the use of healthy, primary human, mouse, and chick muscle cells to generate functional NMJs^21–43^. As a result, species and genetic variations between cell lines and host systems may influence the formation of NMJs. Furthermore, co-culturing motor neurons with healthy skeletal muscles undermines the impact that muscles may have on NMD progression. Thus, co-culturing skeletal muscles and motor neurons generated from the same iPSCs ensures a common genetic background and creates an environment that most mimics the NMD of individual patients.

Previously, in vitro hybrid NMJ models were developed with cells originating from multiple species, using chick or mouse skeletal muscles and co-culturing them with human PSC-derived motor neurons ^33^. This approach is not ideal, since it does not address the need for a fully humanized NMJ system. The models have since evolved to using primary myoblast from human biopsies in co-culture with human PSC-derived motor neurons. Although an advancement from previous methods, this fully humanized NMJ system precludes the investigation of skeletal muscle involvement in NMDs ^34–37^. Finally, recent human NMJ models have been established from PSCs by differentiating these cells into both skeletal muscles and motor neurons in 2D and 3D formats. These previous systems present a remarkable advancement to the studies of human NMJs, however, they require long NMJ formation and maturation time (40 to 60 days), which, restricts their sensitivity and scalability ^42^. Alternative organoid-based models, with shortened NMJ maturation times, although valuable for developmental studies, are often too intrinsically variable to be used for high-throughput disease modeling ^41^.

Thus, in this study, we sought to generate a new 2-dimensional PSCs-derived motor neurons and skeletal muscles co-culture system that spontaneously generates functional NMJs and is amenable to high-throughput screening. Compared to other reported NMJ systems, our protocol is modular, faster, more reproducible, and offers a 6-fold increase in scalability compared to previous models ^40–43^. We characterized our co-cultures morphologically and functionally by immunostaining and optogenetics using a multi well-multielectrode array (MEA) system, respectively. We assayed the sensitivity of our new protocol to effectively model disease-specific NMJ dysfunction using a panel of various PSC lines carrying fALS mutations; *C9orf72* HRE, *SOD1*^A5V^, and *TDP-43*^G298S^. We discovered phenotypic and electrophysiological dysregulation in all ALS-iPSC-derived NMJs compared to controls. The electrophysiological dysregulation observed in NMJ co-cultures of *C9orf72* HRE and *TDP-43*^G298S^, but not in *SOD1*^A5V^, was rescued with GDNF, emphasizing the need for therapeutic studies that accounts for the patients’ genotype. Moreover, the ability of our system to develop functional human NMJs in a reproducible and scalable manner enables the performance of large sample analyses and high-throughput screens designed to uncover new molecular pathways and potential treatments for ALS and other neuromuscular disorders.

## RESULTS

### Human PSC-derived myotubes display canonical morphology and spontaneous contractile abilities

The challenge to generate skeletal muscles from human PSCs has hindered the study of the role of muscle in NMDs. Skeletal muscle is relatively easily reprogrammed from fibroblasts by overexpressing the transcription factor myoblast determination protein 1 (MyoD). However, this strategy does not effectively reprogram stem cells with the same efficiency. Therefore, our lab has optimized a highly efficient skeletal muscle differentiation from PSCs ^44^. We transduced PSCs to stably express MYOD transcription factor and the chromatin remodeling protein BAF60C to drive PSCs into contractile skeletal muscles ^44,45^. We adapted the initial 3D protocol to a 2D system to precisely control the cell number for functional analysis in monoculture and co-culture **(Figure 1A)**. We induced MyoD and BAF60C expression in human PSCs for 72 hours and evaluated the efficiency of myogenesis by immunostaining for the muscle-specific transcription factors MYOD and myogenin (MYOG) on day 3 of differentiation **(Sup.** Figure 1A). We observed that nearly all the cells co-expressed MYOD and MYOG, indicative of myoblast identity **(Sup.** Figure 1B). We also ensured reproducibility in a multi-well format by calculating the standard error mean (SEM) for the expression of MHC (±0.19%, n = 8 replicates) and DES (±0.25%, n = 7 replicates) in our control iPSC-derived skeletal muscles **(Figure 1C)**.

**Figure 1.**
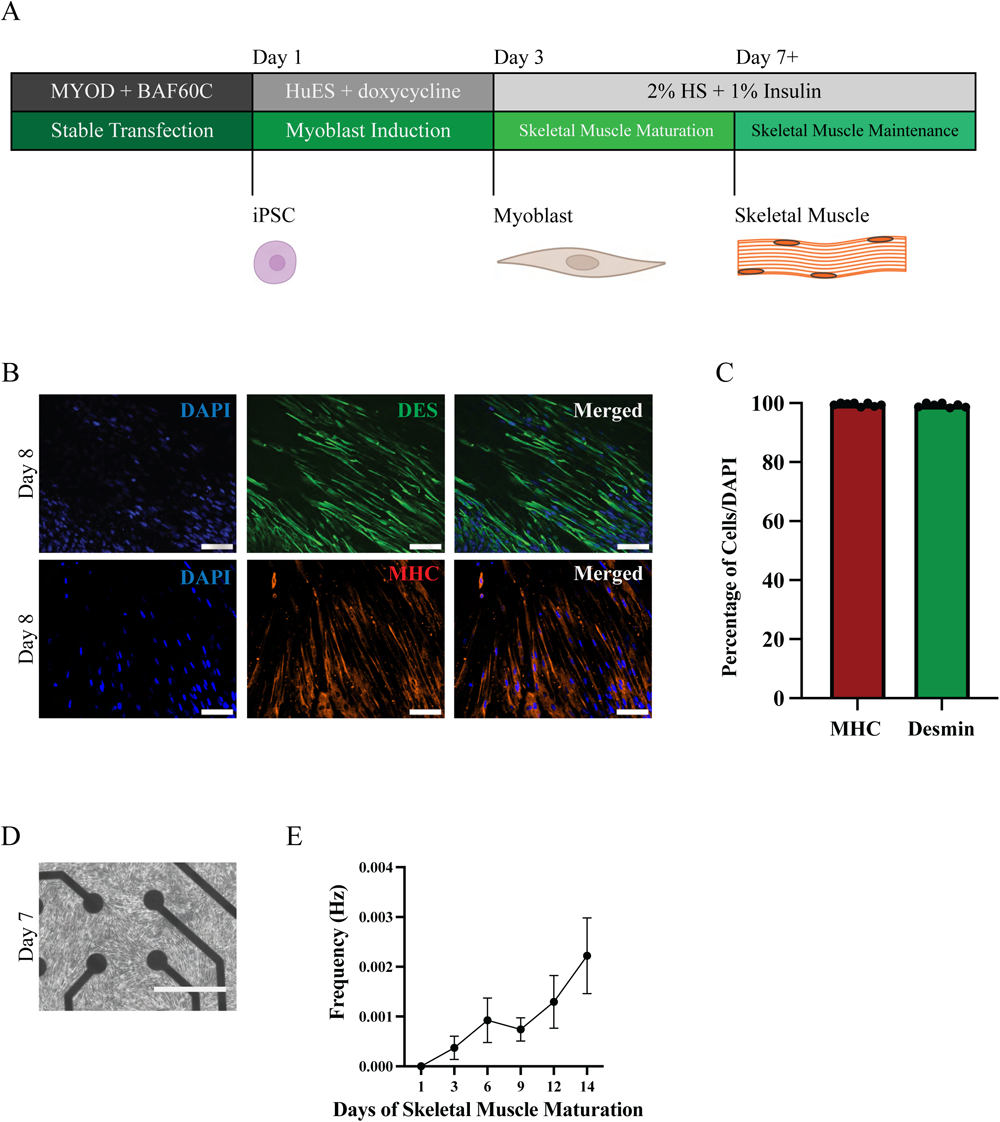
Human PSC-derived skeletal muscles demonstrate markers of maturity and display spontaneous electrophysiological activity. (A) Schematic representation of the differentiation process of hPSCs to skeletal muscles. (B) Immunostaining of hPSC-derived skeletal muscles at day 8 of differentiation for MHC and Desmin, counterstained with DAPI. Scale bar = 100 um. (C) Quantification of the percentage of skeletal muscle cells expressing MHC (4 biological replicates, n = 8 technical replicates per biological replicate) and Desmin (4 biological replicates, n = 7 technical replicates per biological replicate), counterstained with DAPI. Bar graph of average percentage +/- SEM. (D) Brightfield image of skeletal muscles cultured in a well of a multiwell-multielectrode array (MEA) plate. Scale bar = 400 um. (E) MEA recordings of spontaneous skeletal muscle activity frequency (Hz) over 14 days. Each recording was performed for 15 minutes. Line graph of average frequency +/- SEM.

Myoblasts matured into skeletal muscles within one week of differentiation. Fusion of these cells results in a striated appearance of PSC-derived skeletal muscle cultures, resembling the typical morphology of mature skeletal muscles. The PSC-derived skeletal muscles were characterized by the presence of Desmin (DES) and Myosin Heavy Chain (MHC), and as early as day 8 of differentiation nearly 100% of the cells co-expressed these markers **(Figures 1B, 1C)**. We also ensured reproducibility in a multi-well format by calculating the standard error mean (SEM) for the expression of MHC (±0.19%) and DES (±0.25%) in our control iPSC-derived skeletal muscles **(Figure 1C)**. The skeletal muscle cultures also exhibited polynucleation, a typical hallmark of muscle maturation resulting from cell fusion (**Figure 1B)**. Our *C9orf72* HRE, *SOD1*^A5V^, and *TDP43*^G298S^ PSC-derived skeletal muscles were indistinguishable from control PSC-derived skeletal muscles (**Sup.** Figure 1C**, 1D**).

Previous studies have shown that mature skeletal muscles display distinct cellular activities, such as changes in membrane potential and contractility when cultured as myospheres ^41,44^. Thus, we investigated the electrophysiological activity of our optimized PSC-derived skeletal muscle protocol through a Maestro Pro multiwell-multielectrode array (MEA) system to test whether our 2D-cultured skeletal muscles also displayed those characteristics (**Figure 1D)**. For two weeks, we observed measurable membrane potential, confirming that our PSC-derived skeletal muscles are capable of spontaneous activity, which is also seen in 3D muscular cultures. We also noted a gradual increase in this activity, which reflects the progressive maturation of the culture **(Figure 1E)**.

### Human PSC-derived motor neurons display a consistent increase in spontaneous electrophysiological activity and are responsive to optical stimulation

Protocols to differentiate motor neurons from PSCs have been established and substantially optimized since 2008 when the very first disease model using iPSCs was published ^14,46^. Recent 2D and 3D iPSC-NMJ models have shown limited motor neuron differentiation efficiency (5-40%) and wide variability, both within-line and between-line ^40–42^. We chose a 2D spinal cord motor neuron differentiation protocol to enable strict control of motor neuron subtype and number, which would be important when establishing our co-culture. We reasoned that optimization of individual cell type ratios in co-culture would facilitate a reproducible platform for NMJ formation and enable temporal, quantitative measurement of skeletal muscle innervation or de-innervation. Thus, motor neurons were differentiated from healthy human stem cells using a modified version of the dual-SMAD inhibition protocol ^46^ **(Figure 2A)**. To assess differentiation efficiency, we stained for neuronal and motor neuron progenitor (MNP) markers on day 15 of differentiation. Quantification revealed a highly pure MNP population expressing OLIG2, SOX2, and PAX6 **(Sup.** Figure 2A). The PSC-derived motor neuron cultures are mature on day 25 of differentiation (add citation), where 92.49±1.88 and 91.90±1.61 of cells express the motor neuron-specific transcription factors homeobox-9 (HB9) and Islet1/2 (ISL1/2), respectively **(Figures 2B, 2C)**. Furthermore, the cultures display abundant expression of neurofilament H (SMI31), demonstrating the typical neuronal morphology of dendritic arborization. Moreover, there were no GFAP-positive cells, suggesting the absence of astrocytes and non-myelinating Schwann cells **(Figure 2B)**.

**Figure 2.**
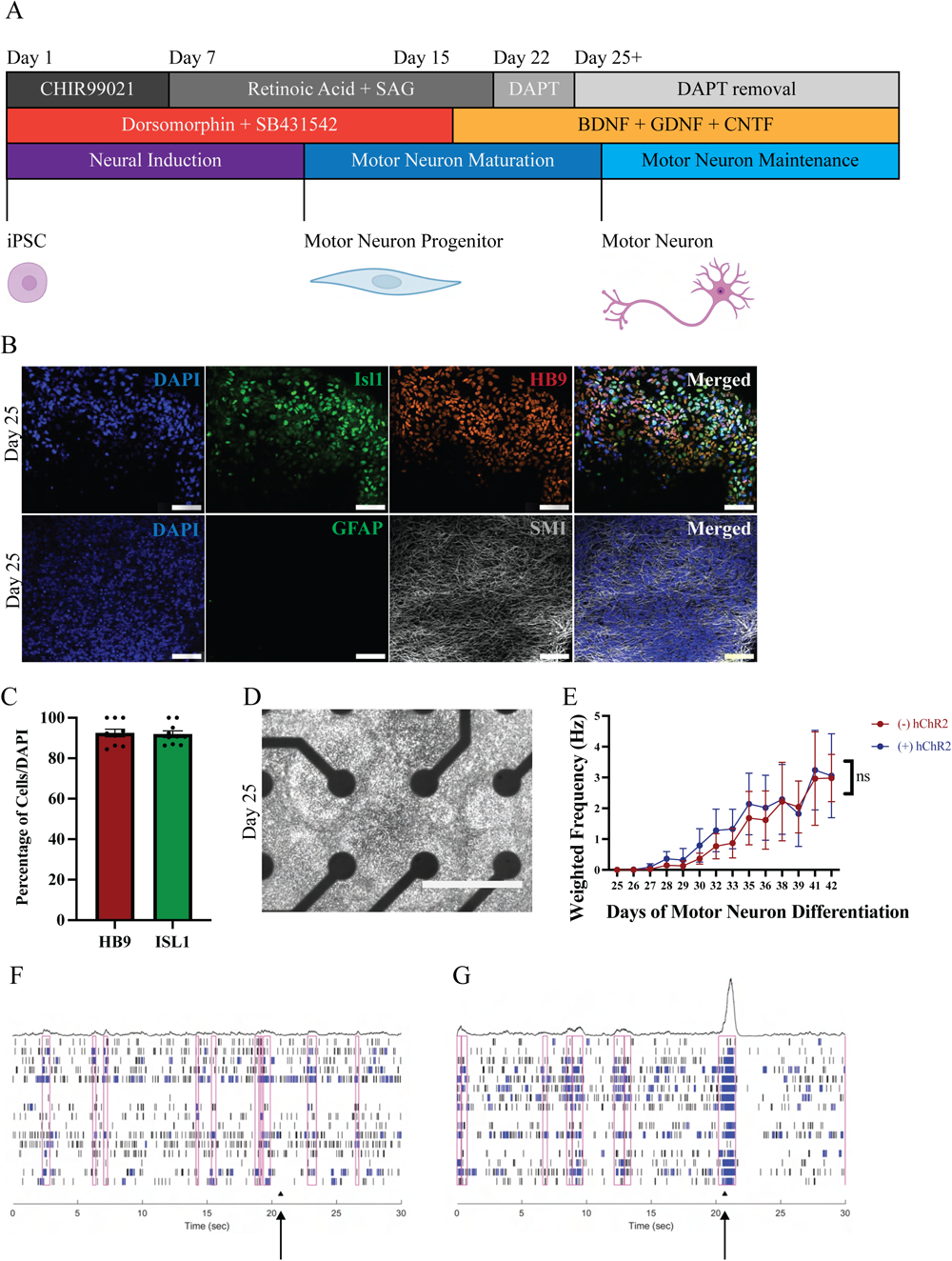
Human PSC-derived motor neurons express markers of maturity and display spontaneous and evoked electrophysiological activity. (A) Schematic representation of the differentiation process from hiPSCs to motor neurons. (B) Immunostaining of motor neuron cultures at day 25 with HB9, Isl1, SMI31, and GFAP, counterstained with DAPI. Scale bar = 100 um (C) Quantification of motor neurons expression HB9 (4 biological replicates, n = 8 technical replicates per biological replicate) and Isl1 (4 biological replicates, n = 8 technical replicates per biological replicate) at day 25 of differentiation. Bar graph of average percentage +/- SEM. (D) Brightfield image of motor neurons cultured in a well of MEA plate. Scale bar = 400 um (E) MEA recording of the weighted mean frequency (HZ) observed in SYP::hChR2-YFP ((+) hChR2) transduced and naive ((-) hChR2) motor neurons over 17 days. Line graph of weighted mean frequency average +/- SEM. ns = not significant. (F) Raster plot of the electrode activity of -hChR2 motor neurons while exposed to blue light stimulation. The arrow represents stimulation. (G) Raster plot of the electrode activity of +hChR2 motor neurons while stimulated by blue light. The arrow represents stimulation.

The PSC-derived motor neurons demonstrated spontaneous action potentials, as measured by multi-electrode array (MEA) recordings, which increased over time as the maturation of the motor neuron cultures progressed **(Figures 2D, 2E).** We tested whether the PSC-derived motor neurons were amenable to optical stimulation by transducing them with the channelrhodopsin-YFP fusion under the control of the synapsin (SYN) promoter (SYN::hChR2-YFP) ^47^. ChR2-transduced motor neuronal activity was measured using the MEA system and compared with non-transduced control PSC-derived motor neurons. Transduced PSC-derived motor neurons showed an increase in burst frequency in response to optical stimulation, without any significant changes to their spontaneous activity compared to non-transduced motor neurons **(Figures 2D-G, Sup.** Figure 2C). Furthermore, we did not observe unstimulated bursts between intervals of optical stimulation **(Figure 2G)**. These findings show that the transduced PSC-derived motor neurons respond to optical stimulation, without altering their intrinsic electrophysiological activity.

### Llower motor neuron density facilitates human PSC-derived NMJ formation

A caveat of translating studies from PSC models to the clinic is the focus on single cell types in detriment to the complex systems of the adult organism. In the motor neuron disease field specifically, PSC investigations have largely focused on preventing motor neuron death. Unfortunately, this approach is no longer sufficient, as prolonging motor neuron survival does not assure reinnervation, nor does it guarantee the prevention of denervation. Thus, we aimed to develop a method to co-culture PSC-derived skeletal muscle and motor neurons that enables spontaneous NMJ formation. To create these 2D human PSC-derived NMJs, we first extensively optimized the dissociation of mature neuronal cultures to improve motor neuron survival and recovery after reseeding. We determined that a 1:1 ratio of Accutase to Accumax effectively separated neurons from their clusters into single cells. Accutase alone lifted neuronal cultures as clumps and required further pipetting to break up the clusters, leading to decreased cell survival.

For NMJ generation, we independently differentiated human PSCs into skeletal muscle and motor neuron cultures. In the motor neuron differentiation process, the notch pathway inhibitor, DAPT, is removed on day 25, the earliest time point when motor neurons co-expressed HB9 and Isl1/2. We dissociated and plated day 25 motor neurons on top of day 5 skeletal muscles **(Figure 3A)**. Prior literature has shown that muscular and synapse maturation occurs when muscle cells are in contact with neurons ^48^. Thus, we plated motor neurons on top of immature skeletal muscles (day 5) to optimize their maturation in direct contact with motor neurons. On day 5 of skeletal muscle differentiation, our muscle cultures no longer express MyoD or myogenin, indicating that they have transitioned out of the myoblast stage and started maturing into myotubes. We then assessed NMJ formation by immunostaining with a nicotinic acetylcholine receptor (nAChR) antagonist, alpha-bungarotoxin (*α*BTX), and its colocalization with synaptophysin (SYP) **(Figure 3B)**. We quantified the number of NMJs on days 3, 5, 7, and 10 (data not shown). When compared to skeletal muscle cultures alone, we observed abundant colocalization of *α*BTX and SYP in co-cultures **(Sup.** Figure 3A). Quantification revealed that the maximum number of NMJs were visualized after 7 days of co-culture. Based upon these encouraging results, we sought to enhance NMJ formation further. Agrin has been described to be essential for the formation and maintenance of NMJs ^49,50^. Therefore, we tested whether the addition of agrin could enhance the expression of nAChR. We found the continuous treatment of agrin (0.1 ug/mL) in our co-cultures increased nAChR protein levels by an average of ∼2.5-fold (±0.22) compared to vehicle control **(Figure 3C, Sup.** Figure 3B). This finding indicates that agrin likely improves human NMJ formation, as well as helps maintain NMJs in our system.

**Figure 3.**
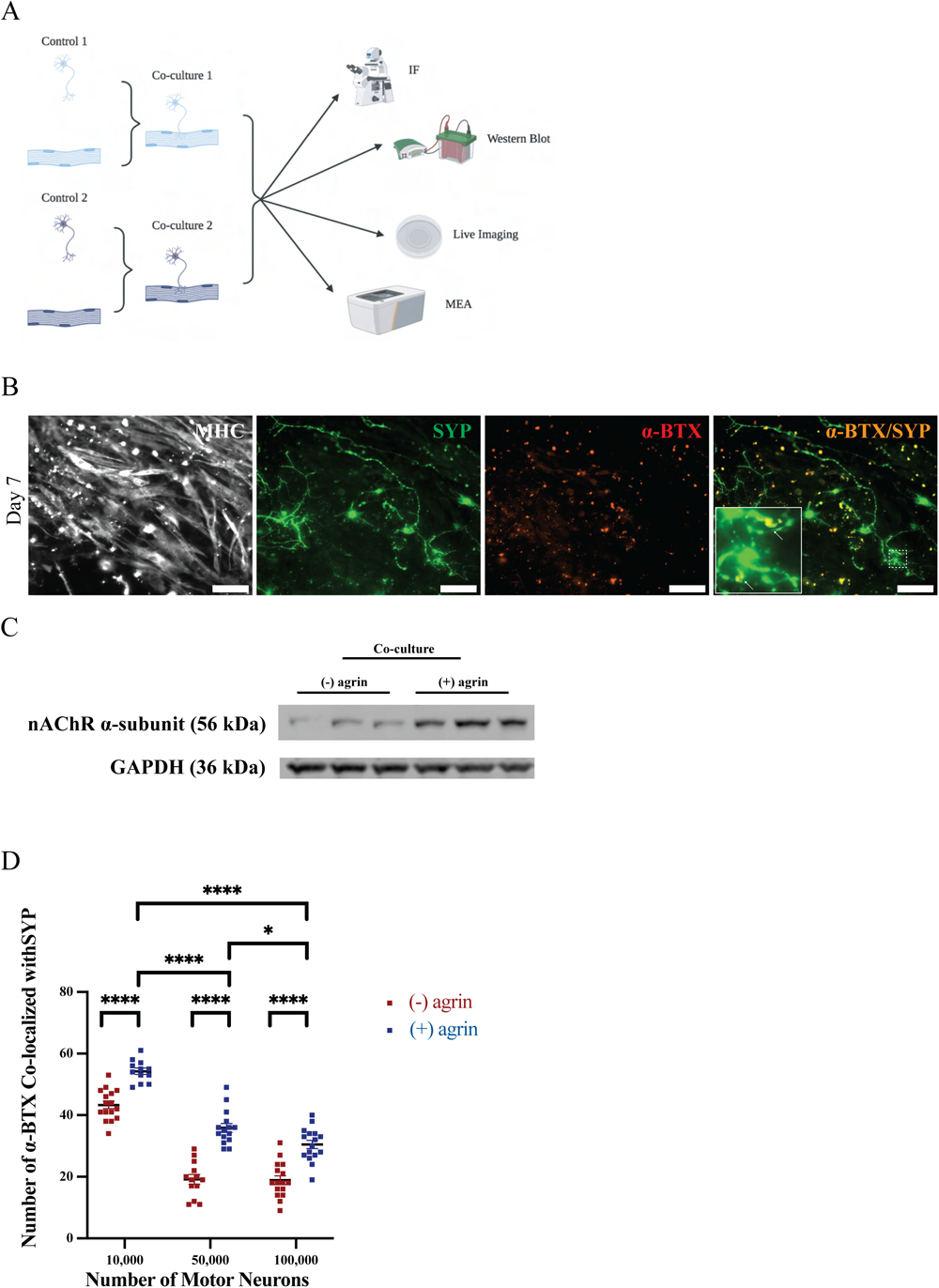
Optimization of human PSC-derived neuromuscular junctions (NMJs). (A) Schematic representation of the experiments to optimize and validate the 2D co-cultures. (B) Immunostaining of motor neurons co-cultured with skeletal muscles. Motor neurons were stained for synaptophysin (SYP, presynaptic site) and muscles were stained for MHC. Synapses were determined by the colocalization of alpha-bungarotoxin (*α*BTX, post-synaptic) and SYP. (C) Western blot for acetylcholine receptor alpha subunit (∼56 kDa) in co-cultures treated with and without agrin. GAPDH (37 kDa) was used as the loading control. (D) NMJ synapse quantification of *α*BTX/SYP co-localization in iPSC-derived co-cultures with 10,000; 50,000; and 100,000 motor neurons with 150,000 skeletal muscles (n_10,000_ = 12, n_50,000_ = 15, and n_100,000_ = 16 technical replicates. Dot plot of the average number of colocalized puncta +/- SEM.

To optimize the motor neuron seeding density to maximize NMJ synapse formation, we compared motor neuron densities of 10,000; 50,000; and 100,000 co-cultured with 200,000 skeletal muscle cells per well (0.7 cm^2^) in the presence and absence of agrin. Quantification of NMJs was assessed based on immunostaining of *α*BTX and SYP co-localization, which revealed between 1.25-and 1.87-fold increase in synapse formation when co-cultures were maintained with agrin, regardless of motor neuron number **(Figure 3D)**. This agreed with our findings of elevated nAChR levels with agrin treatment. Moreover, lower motor neuron seeding density (∼10,000) with skeletal muscles significantly increased the number of synapses formed, when compared to co-cultures with 50,000 or 100,000 motor neurons by 1.51 (±0.06) fold and 1.78 (±0.14) fold, respectively **(Figure 3D)**. Together, these results demonstrate that the presence of agrin improves spontaneous human NMJ formation *in vitro* and lower densities of motor neurons with skeletal muscles are ideal for NMJ formation.

### Human PSC-derived NMJs are functional and optically responsive, and their activity is quantifiable

A bonafide *in vitro* NMJ model should demonstrate the co-localization of pre-and post-synaptic proteins that operate synergistically to form a functional motor unit. Therefore, to assess the functionality of the human PSC-derived NMJs in a quantifiable manner, we optimized spontaneous and optically stimulated electrophysiological readings on an MEA **(Figures 4A, 4B)**. We leveraged the spontaneous electrophysiological activity of PSC-derived skeletal muscles as a read-out for PSC-derived skeletal muscle **(Figures 1D, 1E)** and motor neuron **(Figures 2D, 2E)** derived NMJ functional activity. We observed as much as a 200-fold increase in the weighted mean frequency of PSC-derived skeletal muscle activity when muscles were co-cultured with PSC-derived motor neurons compared to the spontaneous activity of PSC-derived skeletal muscles alone **(Figure 4C, 1E)**. Furthermore, when motor neurons were optogenetically stimulated, there was a significant increase in the weighted burst activity elicited by skeletal muscle **(Figure 4D)**. To show that the electrophysiological signal recorded from the co-culture was a direct result of NMJ activity, we incubated the co-cultures with the nAChR antagonist, curare (60 ug/mL) ^51^. Curare treatment resulted in complete ablation of the electrophysiological signal, confirming that the enhanced activity recorded from the co-cultures originated from the PSC-derived skeletal muscles innervated by the PSC-derived motor neurons **(Figure 4E)**. As expected, there was no significant difference in electrophysiological activity of PSC-derived motor neuron cultures treated with curare, compared to untreated control **(Sup.** Figure 4). This sustained motor neuron activity post-curare treatment demonstrates that the MEA co-culture electrophysiological activity assesses NMJ functionality.

**Figure 4.**
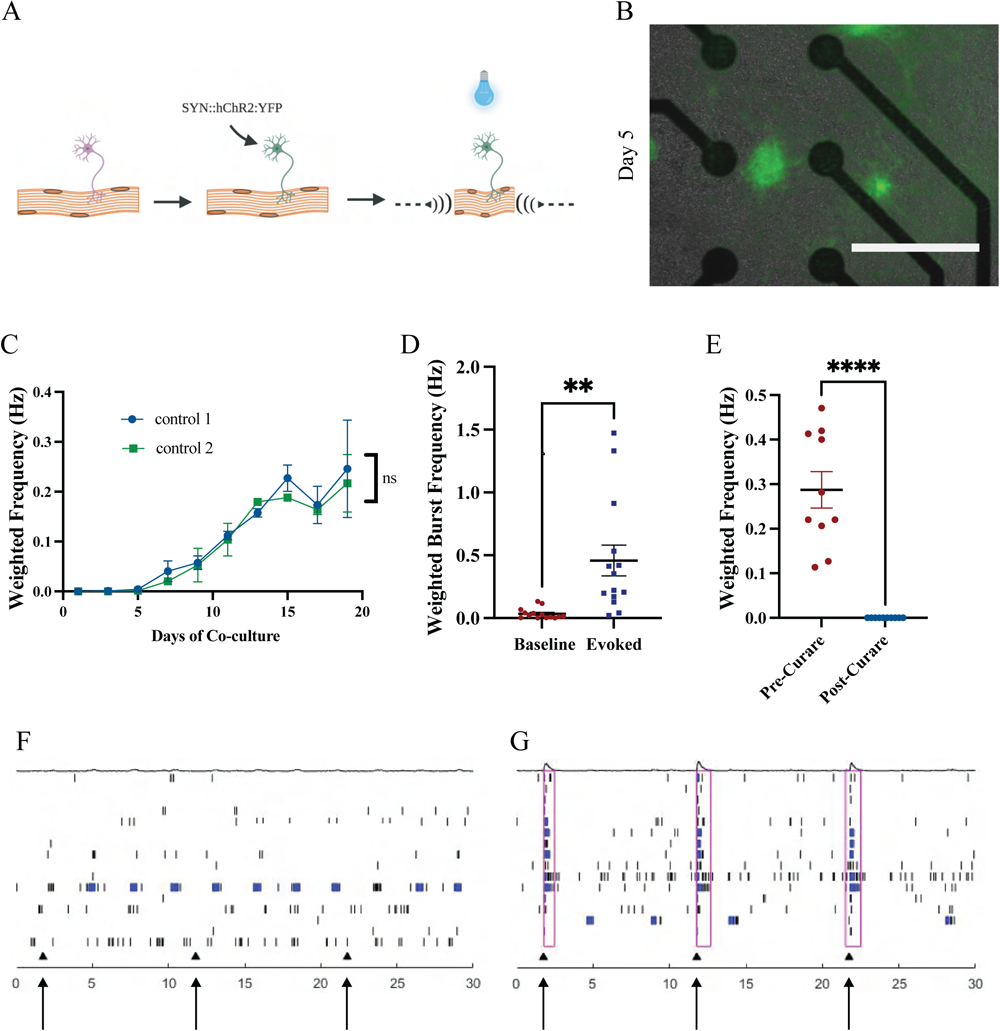
NMJs display consistent electrophysiological activity and can be stimulated with optogenetics. (A) Schematic representation of optogenetic stimulation of (+) hChR2 motor neurons inducing muscle contraction. (B) Image of (+) hChR2 motor neurons co-cultured on top of skeletal muscles in a well of an MEA plate. Scale bar = 400 um (C) MEA recording of the weighted mean frequency of spontaneous co-culture activity (2 biological replicates, n = 12 technical replicates per biological replicate) of two unaffected control lines over 19 days. Line graph of average weighted mean frequency +/- SEM. (D) Comparison of burst activity of muscles in response to baseline and evoked motor neuron stimulation (n = 14 technical replicates). **p<0.01. Dot plot of average burst frequency +/- SEM. (E) Comparison of skeletal muscle innervation before and after curare treatment of co-cultures (2 biological replicates, n = 10 technical replicates per biological replicate). (F) Raster plot of muscle activity in response to (-) hChR2 motor neuron innervation. The arrow represents optical stimulation. (G) Raster plot of muscle activity in response to (+) hChR2 motor neuron innervation. The black arrow represents optical stimulation.

To show the *in vitro* NMJ activity can be directly modulated, we took advantage of optogenetics to stimulate hChR2-transduced PSC-derived motor neuron activity to generate a PSC-derived skeletal muscle response. We initially assessed muscle contraction by live imaging the co-cultures every other day, starting from day 6 of co-culture until there was no more activity observed, ∼ day 18 (± 2) of co-culture. We recorded for 20 seconds without optical stimulation and then 20 seconds with stimulation. As early as 10 days of co-culture, we observed muscle contraction under optical stimulation (∼470 nm) of hChR2-transduced PSC-derived motor neurons co-cultured with PSC-derived skeletal muscles. The PSC-derived skeletal muscle contractions were consistently present until day 16 of co-culture (**Sup. Video 1-8).** The continuously contracting skeletal muscle cultures eventually detached from tissue culture plates, thus our protocol reveals a window of accessibility to study functional human NMJ in a 2D format. To demonstrate reproducibility at scale, we performed the optogenetics-based electrophysiological assessment using MEA with the Lumos apparatus for optical stimulation in medium-throughput 48-well plates. We noted a significant ∼5-fold increase in burst activity in the functional NMJ co-cultures under optical stimulation (Baseline SEM = ±0.01; Evoked SEM = ±0.12) **(Figure 4F, 4G)**. Therefore, our live-imaging results and MEA recordings show that our newly generated human PSC-derived NMJs are functional and responsive to optical stimulation. Furthermore, we demonstrated that we could maintain our co-cultures over an extended time period, which enables temporal analysis of human NMJs in the context of development or disease.

Other studies examining 2D NMJ function have utilized dish or microfluidic devices with compartments for each cell type. By their design, microfluidic devices are restricted by the number of chambers. However, in our established model we measured the synaptic activity of our co-cultures in a 48-well setting, demonstrating at least a 6-fold increase in scale. Moreover, 96-well plates are readily available to the research community, offering a viable path to expand into high-throughput assays. This vastly increases the efficiency of studying NMJ electrophysiology in future experiments.

### *C9orf72* HRE, *SOD1*^A5V^, and TDP43^G298S^ mutations reduce the number of synapses and depress NMJ activity

A research area that would greatly benefit from our human PSC-derived in vitro NMJ method is the NMD field. Previous human NMJ studies have identified synaptic defects associated with NMDs. Those studies have developed inventive methods to quantify NMJ activity, such as video-processing algorithms or the number of contractile sites ^40,43^; unfortunately, those are not scalable. Moreover, the time course for examining NMJ dysfunction only ranges from a few seconds to a few hours. Therefore, there is a lack of research in the NMD field that can consistently study synaptic activity over longer periods of time (days/weeks). Clinical studies and animal models have shown that NMJ loss plays a significant role in ALS disease onset and progression ^3,52^. Thus, we wanted to leverage iPSCs from patients with three different fALS mutations, *C9orf72* HRE, *SOD1*^A5V^, and *TDP43*^G298S^ to test the sensitivity of our new human NMJ protocol to detect synaptic dysfunction in ALS. PSC-derived motor neurons and skeletal muscles were generated from *C9orf72* HRE, *SOD1*^A5V^, and their respective CRISPR/Cas9 engineered isogenic controls, *TDP43*^G298S^ and the widely used embryonic stem cell control line (H9). The human PSC-derived motor neurons and skeletal muscles were then co-cultured as previously described, the number of NMJs was assessed by the colocalization of SYP and *α*BTX **(Figure 5A)**, and the functional analysis of these synapses was assessed using the MEA system **(Figure 6A)**. We reasoned that a comparison of the different CRISPR/Cas9-engineered isogenic controls and the H9 control would show a similar number of synapses and activity due to the robustness and high reproducibility of our new PSC-derived NMJ method. Therefore, we compared the number of synapses and the electrophysiology of the isogenic and H9 control to each other, and we observed no significant differences **(Sup.** Figure 5A).

**Figure 5.**
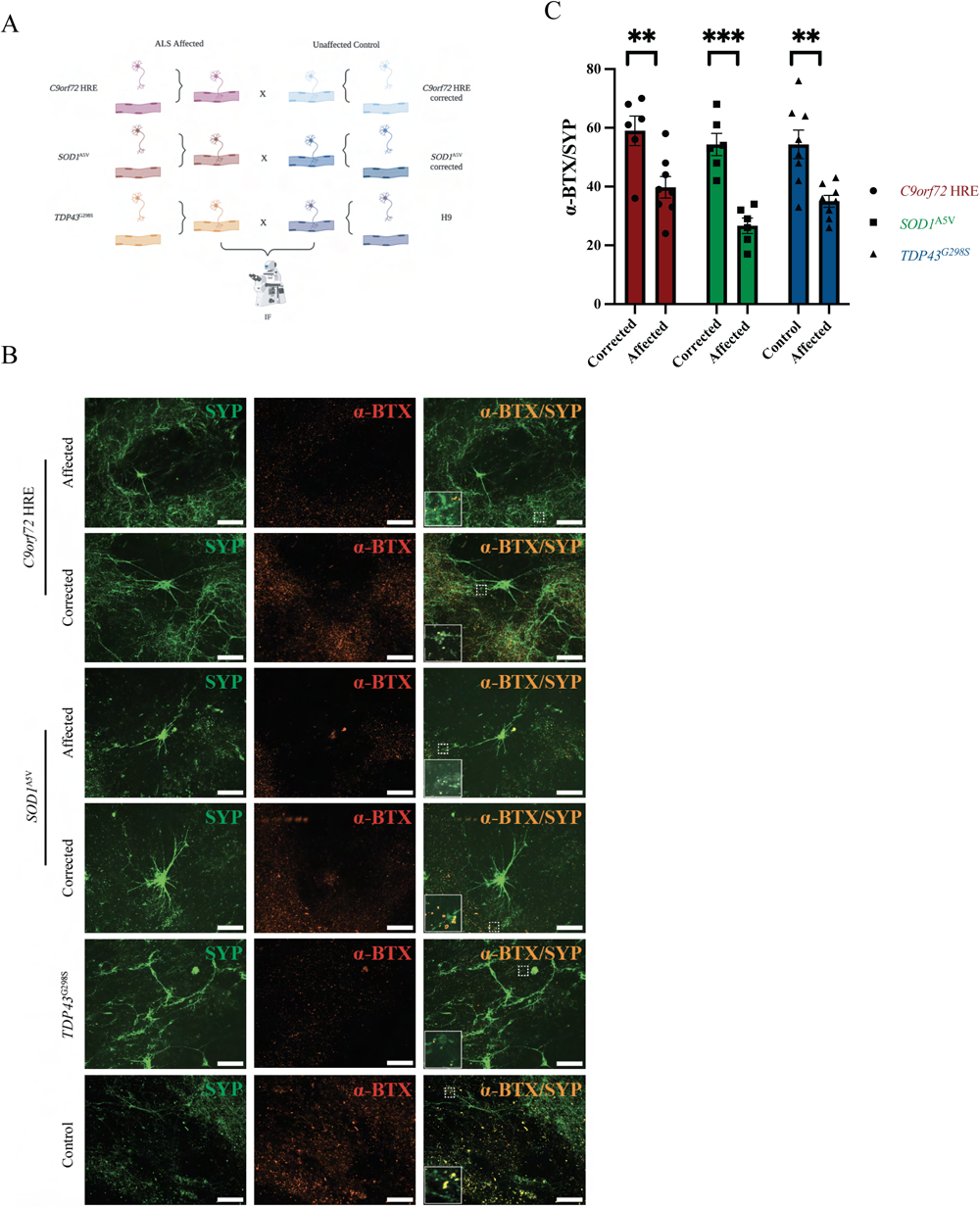
PSC-derived ALS-affected co-cultures show a decreased number of synapses after. (A) Schematic representation of the immunofluorescent comparison of the different ALS mutations and their respective controls (B) Immunostaining of hiPSC-derived co-cultures of *C9orf72* HRE, *C9orf72* HRE corrected, *SOD1*^A5V^, *SOD1*^A5V^ corrected, a TDP43^G298S^, and an unrelated control. (C) NMJ synapse quantification of the different ALS-affected co-cultures with their corrected isogenic pair and an unaffected control.

**Figure 6.**
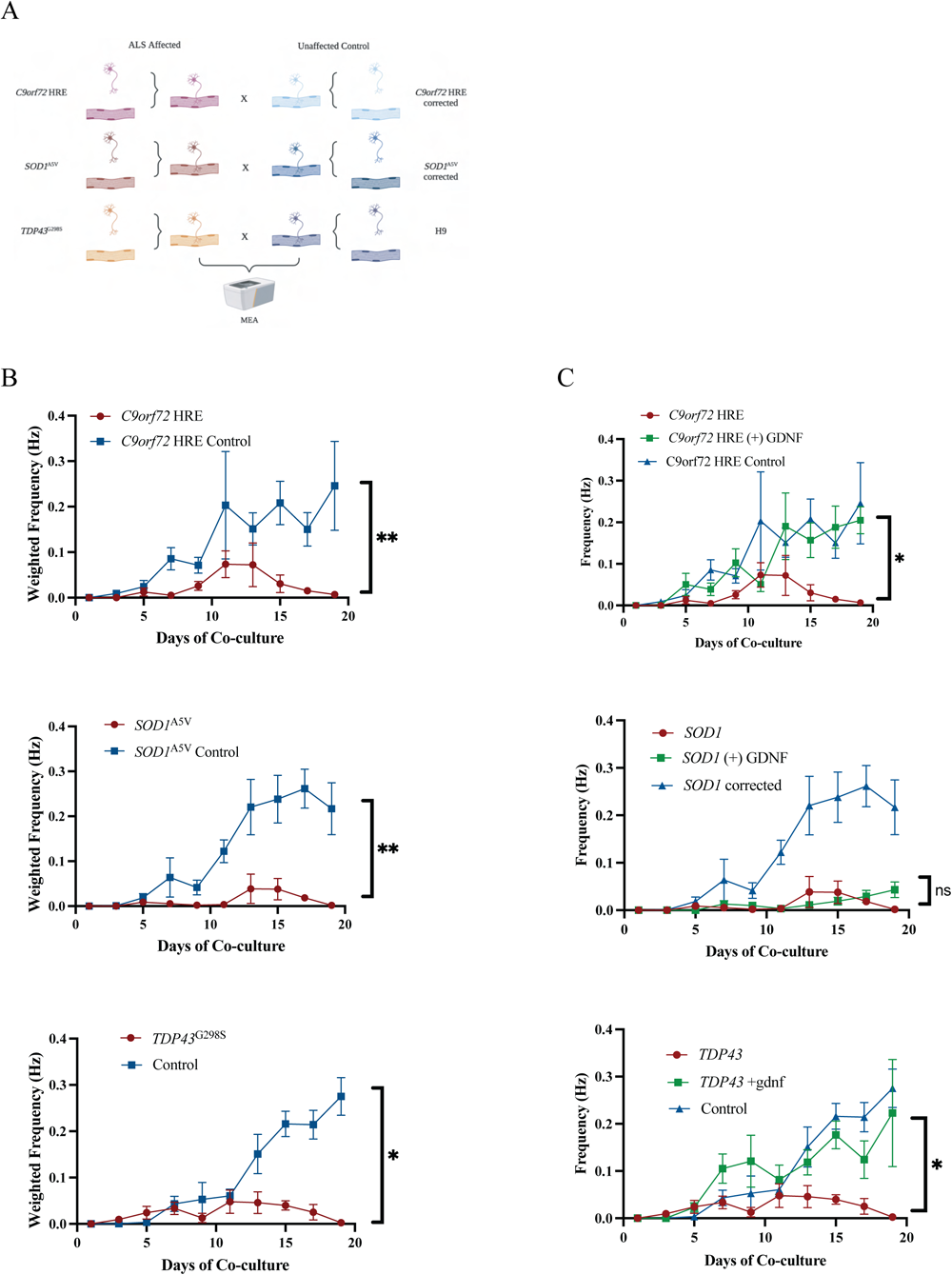
ALS-affected co-cultures display less frequent muscle innervation compared to corrected and unaffected co-cultures. (A) Schematic representation of the electrophysiological comparison of the different ALS mutations and their respective controls (B) MEA recording of the weighted mean frequency of spontaneous co-culture activity (n = 8 technical replicates) of *C9orf72* HRE and *SOD1*^A5V^ with their respective corrected pairs, and a TDP43 mutant with an unaffected control. The electrophysiological activity of the corrected co-cultures was also compared with the unaffected control. Line graph of the average weighted mean frequency +/- SEM. (C) MEA recording of the weighted mean frequency of spontaneous co-culture activity (n = 8 technical replicates) of *C9orf72* HRE and *SOD1*^A5V^, and *TDP43*^G298S^ without GDNF treatment, with GDNF treatment, and their respective control pairs. Line graph of the average weighted mean frequency +/- SEM.

To evaluate the ALS lines for phenotype differences in NMJ function, we next compared the ALS lines to their respective isogenic and H9 controls. The *C9orf72* HRE co-cultures (n=8) showed ∼1.5-fold reduction in the number of synapses formed at day X when compared to the paired isogenic control (n=6) (p*_C9orf72_*=0.0079) **(Figure 5B, 5C)**. The NMJ function in *C9orf72* HRE co-cultures was also significantly reduced (p*_C9orf72_*=0.0066) **(Figure 6B)**. The *SOD1*^A5V^ co-cultures (n=6) displayed ∼2-fold reduction in the number of synapses when compared to its paired isogenic control (n=6) (p*_SOD1_*=0.0001) **(Figure 5B, 5C)**, and similarly to *C9orf72* HRE co-cultures, *SOD1*^A5V^ NMJ activity was also significantly decreased (p*_SOD1_*=0.0056) **(Figure 6B)**. Finally, *TDP43*^G298S^ co-cultures (n=8) displayed ∼1.5-fold fewer co-localized *α*BTX and SYP (n=8) (p*_TDP43_*=0.0026) **(Figure 5B, 5C)** and statistically reduced NMJ activity (p*_TDP43_*=0.031) compared to the H9 control **(Figure 6B)**, in line with the other familial ALS mutations. These results suggest both a morphological and functional defect in ALS NMJs. Our histological assessment and MEA electrophysiological recordings of NMJs derived from ALS patient iPSCs and respective controls recapitulate NMJ phenotypes that have been reported in the literature ^3,40,53,54^. Because of the consistency and scalability of our method, as well as its proven sensitivity to detect relevant disease phenotypes within a week of motor neuron and skeletal muscle co-culture, our platform provides a new, promising avenue for high-throughput screening and potential therapeutic discoveries for ALS and other NMDs.

### GDNF Rescues Synaptic Activity in *C9orf72* HRE and *TDP43*^G298S^, but not in *SOD1*^A5V^

Trophic factors have been considered a promising therapeutic avenue for neurodegenerative diseases. Specifically, glial cell-line derived neurotrophic factor (GDNF) has been known to play a role in neuroprotection and maintaining the neuromuscular synapses^55^. Moreover, a recent phase 1/2a clinical trial that results in the secretion of excess GDNF in the spinal cord of sporadic ALS patients have shown promising results ^56^. After observing the significant reduction in NMJ activity in our 3 different familial ALS cultures, we next wanted to investigate if synaptic communication could be restored. We leveraged our functional assay using the MEA to test if GDNF treatment could rescue the synaptic defects identified in our ALS iPSC co-cultures. Thus, the co-cultures were treated with 20ng/ml of GDNF for X days and assessed for increased electrophysiological activity. We observed a restoration of NMJ synaptic activity in the *C9orf72* HRE (n=8) (p*_C9orf72_* _(+) GDNF_=0.0136) and *TDP43*^G298^ co-cultures (n=8) (p_TDP (+) GDNF_ = 0.0108) (**Figure 6C**). However, GDNF treatment of *SOD1*^A5V^ co-cultures did not result in any significant differences compared to the untreated co-cultures (n=8) (p_SOD1_ = 0.8353) (**Figure 6C**). Collectively, the administration of GDNF and the genotype dependent outcomes in the 3 ALS mutations supports the paradigm of mutation specific therapeutic intervention.

## DISCUSSION

The complex processes of initiation and coordination of signaling between the central nervous system (CNS) and muscles are highly susceptible to damage in aging, developmental disorders, and NMDs ^7,57^. This vulnerability is positioned most prominently at the NMJ, where motor neurons generate muscle contraction by releasing acetylcholine into the synaptic cleft. Sadly, partially due to a lack of complete scientific understanding of the human NMJ, currently, there is a lack of effective treatment for most NMDs characterized by dysfunctional NMJs. Early studies on synaptogenesis and motor neuron-to-skeletal muscle communication relied primarily on animal models ^9,58^. Eventually, cross-species, particularly human-mice, co-culture systems were developed ^59^. Consequently, while the cellular and molecular features of mouse NMJs have been characterized in depth, there is much less known about human NMJ. Thus, it is unsurprising that drugs found to alleviate symptoms in NMD animal models have frequently failed in clinical trials ^60,61^. In order to expand our understanding in ways that will lead to effective treatments for patients, the field needs to move beyond animal models and directly study human NMJs. The iPSC technology allows for the genomes of patients afflicted with NMJ impairment to be captured in a pluripotent cell line. The investigation of NMD pathology has been performed using affected cells, such as motor neurons, and skeletal muscles, for example, differentiated from PSCs. Therefore, here we present a new human stem cell derived NMJ model that is highly reproducible across different lines, responsive to optogenetics, and suitable for high throughput screening.

Prior investigations have highlighted the challenges in establishing 2D NMJs from a single iPSC ^62^. Some of these challenges include difficulty discerning the different cell types and maintaining cell cultures over time, given the natural contractility of skeletal muscles, which ultimately leads to their detachments from the culture dish ^35^. Concerns regarding the maintenance of 2D models, especially the viability of contracting skeletal muscles, led to research towards other methods, such as spheroid development ^38,40,41^. 3D NMJ systems can be advantageous when it comes to depicting how neurons, glial, and other cell types interact with each other over extended periods of time. Recent models have been able to generate self-organizing organoids, sensorimotor organoids, and corticomotor assembloids that resemble the motor cortex and spinal cord ^39–41^. These models are promising for studying developmental processes; however, one major caveat with organoid modeling is their reproducibility ^63^. Recent studies have shown improvements in this area; however, there is still considerable variability in both intra-and inter-cell lines ^39,40^. Given this variability, we pursued an NMJ model using a 2D approach to effectively control the seeding number of motor neurons and skeletal muscle and ensure the experimental sensitivity of the system. Our PSC-derived motor neuron and PSC-derived skeletal muscle 2D co-culture protocol also generate spontaneous functional NMJs up to 10x quicker than previously published protocol ^42^. Moreover, we engineered our co-culture system to be modular. By differentiating PSCs into motor neurons and skeletal muscles separately, our protocol enables the co-culture of cells derived from affected and unaffected individuals. This gives our model the unique advantage of investigating the contribution of different cell types to disease pathogenesis (i.e. NMD patient iPSC-derived motor neurons co-cultured with unaffected iPSC-derived skeletal muscle and vice-versa).

Functional NMJs in a 2D culture have been previously generated, albeit it requires over 60 days for cell differentiation and maturation. Furthermore, this method has also described the presence of undesired cell types, such as different neuronal subtypes and glia ^42^. While glia may play an important role in the development and maintenance of NMJs, we deliberately developed a system to interrogate motor neuron and skeletal muscle interactions. If desired, additional cell types could be co-cultured in a controlled manner. Here, we leveraged protocols that promote human PSCs to efficiently differentiate into homogenous cultures of motor neurons and skeletal muscles, separately. As shown in the skeletal muscle PSC-derived cultures nearly 100% of the muscle cells expressed MHC **(Figure 1)** and in the PSC-derived motor neuron cultures, over 90% of the cells expressed HB9 **(Figure 2)**. Their co-culture, then allowed for spontaneous human NMJs to be formed within a few days (∼5 days) **(Figure 3)**. Furthermore, we determined the optimal number of PSC-derived motor neurons to be co-cultured with PSC-derived skeletal muscles to maximize the number of functional synapses. Here, we saw that more synapses were formed when motor neurons were plated at a lower density (Figure 3D). We hypothesize that this occurred due to a potential preference of neurons to form connections with other neurons rather than alternative cell types. Therefore, by decreasing the number of PSC-derived motor neurons distributed onto the PSC-derived skeletal muscles, we limited the number of neuron-to-neuron synapses and favored connections to the skeletal muscle. Another challenge to the long-term co-culture of PSC-derived motor neurons and PSC-derived skeletal muscles, and the efficient development of stable NMJs, is the maintenance of skeletal muscles in 2D culture. As they mature, spontaneous contractions occur more frequently, and the cultures eventually detach from the plate surface ^64^. Therefore, the rapid NMJ formation described here (∼5 days of co-culture) allows for the co-cultures to remain stable and attached to the plates for a longer period of time (∼18 days) than previously reported ^64^.

The multi well-multielectrode array system we used in this study provides an innovative approach to measure the electrophysiology of PSC-derived skeletal muscle, motor neurons, and co-cultures over time. We were able to quantifiably record spontaneous PSC-derived skeletal muscle activity that gradually increased over time using an MEA system. Although the number of spikes recorded from our muscle monocultures was limited, this is a reflection of absent stimulation. Our optimization of the use the MEA system to measure activity from skeletal muscles, enabled this system to analyze functional NMJs *in vitro*, even in the absence of visible muscle contractions. In co-cultures, the electrophysiological activity of our PSC-derived skeletal muscle drastically increased upon PSC-derived motor neuron innervation.

The activity of unaffected control skeletal muscles continued to be present over ∼15 days of co-culture. Generating monocultures and co-cultures on top of electrodes, in a multi well format also enabled us to apply specific treatments to individual wells and observe their effects in real time, which may not be possible with traditional patch-clamping methods. By applying optogenetics, we were able to induce more robust contractions in PSC-derived skeletal muscles. Optogenetics is a relatively new and refined system of controlling cellular functions ^65^. Upon optical stimulation, photosensitive proteins mediate changes in cell membrane potential and elicit action potential ^65,66^. Previous studies have shown myosphere contraction when cultured with optogenetically stimulated spinal cord motor neurons ^40,41^. In our study, however, we applied the same optogenetic principles and stimulated PSC-derived motor neurons to elicit a controlled and robust muscular contraction which then could be recorded by an MEA system and visualized by live imaging. We were able to extend our culture for ∼18 days, despite consistent spontaneous muscle contractions. This window of time will be crucial for future longitudinal experiments.

Our PSC-derived NMJ model provides an important advancement for therapeutic investigation for various NMDs, such as, but not only limited to, ALS. For decades the ALS field has benefited from PSC-derived motor neurons to understand and identify novel intracellular mechanisms associated with ALS pathogenesis. However, a general caveat of studying ALS, or other NMDs, with iPSC models has been the focus on individual cell types while overlooking complex multicellular systems present in organisms. In the NMD field specifically, iPSC investigations have largely focused on addressing phenotypes that could prevent motor neuron degeneration ^67^. However, prolonging motor neuron survival does not guarantee muscle re-innervation ^68,69^. Our PSC-derived NMJ model presents an opportunity to focus on identifying cell-specific targets and testing those targets on the maintenance and sustained human motor neuron innervation of human skeletal muscles. Therefore, to demonstrate the disease modeling potential of our system, we performed morphological and functional NMJ-specific assessments using three different familial ALS-derived iPSCs, including C9orf72 HRE, *SOD1*^A5V^, and *TDP43*^G298S^.

Zebrafish and mouse models have been useful to elucidate ALS pathogenesis associated to C9orf72 HRE. Different mouse models have been developed to study C9orf72 HRE, however the NMJ and motor neuron phenotypic presentation are variable ^70^. While motor neuron abnormalities, axon degeneration and NMJ deinervation are present in a BAC transgenic mouse model containing the full-length C9orf72 gene but it’s absent in most other models ^71–76^. In contrast, a reduced number of acetylcholine receptors at NMJs and motor behavioral deficits have been well-described in the zebrafish model ^53^. Moreover, iPSC modeling of this familial ALS mutation has been limited. Our iPSC-derived model agrees with prior zebrafish model presenting a reduction in the number of synapses and defective synaptic activity **(Figures 5C, 6B)**. Previous *SOD1*^G93A^ mice studies have described NMJ denervation, even before the onset of motor symptoms ^3,77,78^. Studies that examined the functionality of Soleus muscles in *SOD1*^G93A^ showed significant decreases in contractile kinetics and specific forces due to NMJ alterations ^79^. Furthermore, Pereira et al. were able to model mutant-*SOD1*^G85R^ NMJ using iPSC-derived sensoriorganoids. Their work highlighted changes in the percentage of NMJ innervation and the area of innervated NMJs ^40^. Our quantification of *SOD1*^A5V^ iPSC-derived NMJs shows a comparable reduction in the number of intact NMJs; furthermore, we also identified a decrease in NMJ-specific function **(Figures 5C, 6B)**. Pereira et al. also interrogated the effects of *TDP43*^G293S^ on human-derived NMJs as well. The TDP43-affected sensoriorganoids showed a slight decrease in the average percentage of innervated NMJs, but no significant difference when compared to an isogenic control ^40^. Interestingly, our *TDP43*^G298S^ human PSC-derived NMJs model display a significant synaptic decrease in the number and function when compared to a healthy control **(Figures 5C, 6B)**. We find that our human PSC-derived NMJ system can model familial ALS pathogenesis in an efficient and robust manner when previously published protocols, thus providing a more sensitive platform to enable the identification of NMJ-specific dysfunction in different NMDs.

Furthermore, we also identified GDNF as a potential therapy for C9orf72 and TDP43-mediated forms of ALS in a follow up to a phase 1/2a clinical trial ^56^. Contrarily, GDNF did not appear to have any significant effects of restoring NMJ function in SOD1-ALS. Our findings in our iPSC-NMJ model agree with previous rat studies where despite preserving motor neuron death, supplemental GDNF treatment did not maintain neuromuscular connections or restore functionality ^80^. We speculate that this is a result of different mechanisms that these ALS-causing mutations have on both motor neurons and skeletal muscle.

Thus, supplementary studies need to be conducted to further elucidate the differences in the pathophysiology of different ALS mutations. Additionally, we anticipate that this new technology will be a valuable tool for high-throughput screening to identify novel neurotherapeutics in the future.

## METHODS

### Ethics Statement

All experiments were carried out according to the human stem cell (hSCRO) protocol approved by Case Western Reserve University (CWRU)

### Culturing human induced pluripotent stem cells

The unaffected human PSCs used in this study were generated from a healthy donor and previously published and validated ^81^. Cultures were tested bi-weekly for mycoplasma and maintained in an antibiotic-free medium. The PSCs were maintained on 6 cm cell culture plates (Corning) coated with vitronectin (VTN-N) (Thermo Fisher; 5 ug/mL) in StemFlex medium (Thermo Fisher). To pass the PSC colonies for maintenance and expansion, 1 mL ReLeSR dissociation reagent (Stemcell Technologies) was added to the culture plate for 30 seconds at room temperature, then the ReLeSR was aspirated until a thin film remained on top of the cells. Cells were then incubated for 3-5 minutes at 37 degrees Celsius. Cells were then gently lifted off and resuspended in StemFlex medium and distributed to new 6 cm plates.

The *C9orf72* HRE, *SOD1*^A5V^, *TDP43*^G298S^, and the CRISPR-corrected cell lines were obtained from the Cedars-Sinai Biomanufacturing Center for iPSC lines **(Table 1)**.

### Generating skeletal muscles from transfected human PSCs

We used the Neon Transfection System for creating cell lines with the MyoD, BAF60C, and Helper constructs. Cells were dissociated with Accutase (Thermo Fisher) and passed through a 40 uM mesh. 4 ug of MyoD, 4 ug of BAF60C, and 2 ug of Helper constructs ^44^ were electroporated into 1,000,000 cells at 1400V/20ms/2 pulses.

Before inducing myogenesis in the cells, we first selected the cells that were efficiently expressing the MyoD and BAF60C constructs. We treated the cells with Puromycin (Thermo Fisher; 1.5 ug/mL) and Blasticidin (Thermo Fisher; 3 ug/mL) for 48 hours. After treatment, the cells were given a full media change and 1-2 days to recover. At around 70-80% confluence, myogenesis was induced by switching the medium to HuES media without FGF and with doxycycline (Sigma; 200 ng/mL). Cells were cultured in this medium for 2 days. Myoblasts were then lifted from the plate using Accutase (ThermoFisher) and plated onto surfaces coated with Vitronectin. After plating, the cells were kept in HuES media with doxycycline and Y-27632 (Peprotech; 5 uM) for another day. The following day, the medium was changed to a media with DMEM/F12 + 2% Horse Serum (Sigma) + 1% Insulin Transferrin Selenium (ITS) (Corning) with Y-27632 (5 uM) (Skeletal muscle maturation media) **(Sup.** Figure 5B).

HuES media is made of: 158 mL of DMEM/F12 (Life Technologies), 40 mL of KSR (Life Technologies), 2 mL of non-essential amino acids (NEAA) (Gibco), and 400 uL of beta-mercaptoethanol (Life Technologies).

### Generating motor neurons from human PSCs

Motor neurons were created from a modified version of the dual-Smad inhibition protocol ^46^. PSCs were cultured on 6 cm treated plates (Genesee Scientific) until they reached about 75% confluence. The base medium used for motor neuron differentiation was DMEM/F12 with 0.5x N2 (Gibco), 0.5x B27 (Gibco), PenStrep (Thermo Fisher; 100 ug/mL), and ascorbic acid (Millipore; 0.2 mM). This is referred to as motor neuron media. On days 1-6 of motor neuron differentiation, the culture medium was changed to motor neuron media with dorsomorphin (TOCRIS; 1 uM), SB 431542 (Millipore; 10 uM), CHIR99021 (Peprotech; 3 uM), and Y-27632 (Peprotech; 5 uM). On day 7, cells were split 1:3 from 6 cm plates to 10 cm plates (Gennesse Scientific). From day 7 to day 15, cells were maintained in the motor neuron media with dorsomorphin (1 uM), SB 431542 (10 uM), Retinoic Acid (Millipore; 1.5 uM), SAG (Sigma; 200 nM), and Y-27632 (10 uM). On day 15 of differentiation, cells were dissociated with Accutase and replated in 6-well plates (Corning) coated with Poly-Ornithine (Millipore; 10 ug/mL) and Laminin (Invitrogen; 2.5 ug/mL). 500,000 cells were plated per well and cultured in motor neuron media with BDNF (2 ng/mL), CNTF (2 ng/mL), GDNF (2 ng/mL), Retinoic Acid (1.5 uM), SAG (200 nM), and Y-27632 (5 uM). Cells were left in this medium until day 21. On day 22, the media was changed to the motor neuron media with BDNF (2 ng/mL), CNTF (2 ng/mL), GDNF (2 ng/mL), DAPT (2 uM), and Y-27632 (5 uM). DAPT was removed on day 25 of differentiation. On day 25, motor neurons can be dissociated and replated, frozen, or maintained with media changes every 1-2 days **(Sup.** Figure 5B).

Motor neurons were dissociated using a 1:1 dissociation solution of Accutase and Accumax (Innovative Cell Technologies Inc.) were counted using Trypan Blue to ensure a survival rate above 80%. Cells were incubated in the dissociation solution for 10-15 min at 37 degrees Celsius. After the time had elapsed, the plates were tapped against a surface to lift the cells. Cells were then triturated a maximum of 10 times and passed through a 70 uM strainer. It was important to minimize the number of triturations to maximize cell viability. Cells were then centrifuged at 200 xg for 3 min. Motor neurons and motor neuron progenitor cells can also be plated onto glass surfaces coated with Poly-Ornithine (100 ug/mL) and Laminin (5 ug/mL). Motor neurons could also be frozen in homemade freezing media. For 50 mL of motor neuron freezing media: 25 mL of motor neuron media, 20 mL of FBS (Gemini), 5 mL of DMSO, and 10 uM of Y-27632.

### Transducing Motor Neurons with hChR2

We created lentiviruses using PEI transfection of Lenti-X HEK-293 cells (Clonetech). HEK-293 cells were cultured on 15 cm plates with DMEM with 4.5 g/L of glucose + 10% FBS (HEK media) until they reached about 50% confluence. For every 12 plates of HEK-293 cells, 500 uL of blank DMEM was mixed with SYN::hChR2:YFP (12.2 ugs), pMDLg/pRRE (addgene #12251) (8.1 ugs), pRSV-REV (addgene #12253) (3.1 ugs), pCMV-VSV-G (addgene #8454) (4.1 ugs), and PEI (Fisher; 1 ug/uL). The solution was incubated with HEK-293 cells overnight at 37 degrees Celsius, then the medium was replaced with fresh HEK media and incubated for 3 days. HEK medium was then collected after 3 days and centrifuged at 500 xg for 10 minutes. The supernatant was transferred to a fresh conical tube and the Lenti-X concentrator was added per manufacturer instructions (Clonetech). The supernatant was centrifuged at 1500 xg for 45 minutes or until the viral pellet became visible. The supernatant was then removed and the viral pellet was resuspended in 500 uL of DMEM/F12. 100 uL of the viral suspension is added per 1-2 million cells. The virus can be added between day 15 and day 25 of motor neuron differentiation. We observed strong expression of the hChR2 construct approximately 4-7 days following transduction, as yellow or green fluorescence.

### Creating two-dimensional neuromuscular junctions (NMJs)

Myoblasts were plated in a 48-well clear-bottom plate coated with vitronectin at a cell concentration of ∼150,000 cells per well. Myoblasts were allowed to mature in skeletal muscle maturation media (DMEM/F12 + 2% Horse Serum + 1% ITS) for 2 days. Mature motor neurons were then plated on top of the myotubes at a concentration of ∼10,000 per well. The co-culture was then allowed to mature with careful media changes every other day. Co-culture media is composed of DMEM/F12 + 2% HS + 1% ITS with BDNF (2 ng/mL), CNTF (2 ng/mL), GDNF (2 ng/mL), and Y-27632 (5 uM). For the clustering of NMJs, agrin (R&D Systems; 0.1 ug/mL) was added **(Sup.** Figure 5B).

NMJ experiments in the ALS co-cultures, excluded agrin supplementation to prevent potential masking of NMJ phenotypes. GDNF supplementation, for the rescue experiments, was added at 10x the basal concentration (20 ng/mL).

### Recording Skeletal Muscle, Motor Neuron, Co-culture, and Stimulated Activity

For the recording of the electrophysiological activity of skeletal muscles, the surface of an MEA-plate was coated with vitronectin (Thermo Fisher, 1:50 dilution). Myoblasts were plated after 2 days of doxycycline addition, at a density of 200,000 cells per well of a 48-well plate with 16 electrodes. Media changes were performed before the recordings. Each recording was done for 15 minutes at 37 degrees Celsius and 5% CO_2_. For the recording of the electrophysiological activity of motor neurons, the surface of an MEA plate was coated with poly-ornithine (100 ug/mL) and laminin (5 ug/mL). Motor neurons on day 25 of differentiation were plated at a density of 50,000 per well of a 48-well plate. Media changes were done before the recordings. Each recording was done for 5 minutes at 37 degrees Celsius and 5% CO_2_. For co-culture recordings, myoblasts were plated and allowed to mature for 5 days. Motor neurons were plated on top of the myoblasts. We determined the optimal seeding number to be 10,000 motor neurons on 200,000 skeletal muscles in a 48-well MEA plate. A full media change was done 48 hours later. Full media changes were done 15 minutes before recording. Each recording was done for 5 minutes at 37 degrees Celsius and 5% CO_2_. We stimulated the activity of motor neuron cultures and co-cultures that had been successfully transduced with the channelrhodopsin lentivirus by using the Lumos component of the MEA system using the blue light option. Recordings with and without stimulation were collected. For our purposes, we stimulated our motor neuron cultures every 45 seconds at 50% intensity for 5 minutes and our co-cultures were stimulated every 15 seconds at 50% intensity for 5 minutes. Each recording was collected with the Maestro MEA system (Axion Biosystems) using the AxIS Software Spontaneous Neural Configuration for spontaneous activity and the AxIS Software Optically Evoked Configuration when cultures were stimulated with the Lumos. Axion Biosystems’s Neural Metrics Tool classifies electrodes with at least five spikes per minute as active. Bursts were identified using an interspike interval (ISI) threshold requiring a 5-spike minimum and 100-ms maximum ISI.

### Statistical Analysis of MEA recordings

Transduced and non transduced motor neuron activity and the burst activity of spontaneous and evoked co-cultures were compared by an unpaired Student’s t-test, assuming both populations had the same standard deviations. Analysis of curare treatment of co-cultures and motor neuron populations was analyzed with a paired parametric t-test **(Sup.** Figure 5B). For all statistical tests, a value of *p* ≤ 0.05 was considered statistically significant, with *p* ≤ 0.05 denoted as *, *p* ≤ 0.01 denoted as **, *p* ≤ 0.001 denoted as ***, *p* ≤ 0.0001 denoted as ****, and ns is not significant.

### Immunostaining of human iPCS-derived skeletal muscles and motor neurons

Myoblasts were stained 3 days after induction of differentiation. Skeletal muscles were stained after a week of differentiation. Motor neuron progenitors were stained on day 15 of differentiation. Motor neurons were stained on day 25 of differentiation. The co-cultures were stained at least two weeks after motor neurons were cultured with the skeletal muscles. Cells were first washed with 1xPBS for 5 minutes and fixed with 4% paraformaldehyde (PFA) for 12-15 minutes. PFA was then removed, and cells were washed with 1xPBS 2 times for 5 minutes. Cell membranes were permeabilized with 0.25% triton-x100 in 1xPBS for 10-15 minutes. 0.25% triton-x100 was removed and cells were washed with 1xPBS for 5 minutes. Blocking was performed by adding 3% BSA in 1xPBS for at least 4 hours at 4 degrees Celsius. After blocking, primary antibodies were added and incubated overnight at 4 degrees Celsius. The following day, primaries were removed, and the corresponding secondary antibodies were added for an hour at room temperature. After 1-hour, secondary antibodies were removed and Hoescht 33342 (Thermo Fisher) was added for 5-10 minutes at room temperature. Hoescht was diluted at 1:10,000 in 1xPBS. Then the Hoechst solution was removed, and cells were washed with 1xPBS 3 times for 5 minutes. Cells were mounted with Vectashield (Vector Labs, Inc) to prevent loss of fluorescence, and imaged with an inverted microscope from Zeiss AxioObserver with Apotome **(Sup.** Figure 5B).

The primary antibodies used for myoblasts were mouse anti-myogenin (DSHB; 1:500 dilution) and rabbit anti-MyoD (Santa Cruz Biotechnology; 1:500 dilution). The primary antibodies used for skeletal muscles were mouse anti-MHC (DSHB; 1:500 dilution), and mouse anti-Desmin (Agilent; 1:500 dilution). All dilutions were done in 1xPBS.

The primary antibodies used for motor neuron progenitors were goat anti-OLIG2 (R&D Systems; 1:1000 dilution), mouse anti-nestin (Abcam; 1:1000 dilution), rabbit anti-PAX6 (Biolegend; 1:500 dilution), and mouse anti-SOX2 (Thermo Fisher; 1:500 dilution). The primary antibodies used for motor neurons were: SMI31 (Millipore Sigma; 1:1000 dilution), HB9 (DSHB; 1:1000 dilution), and ISL1 (DSHB; 1:1000 dilution). All dilutions were done in 1xPBS. The HB9 and ISL1/2 antibodies were pre-conjugated with Alexa fluor 555 and 488, respectively, using Antibody Labeling Kit (Thermo Fisher).

The secondary antibodies that were used were: Alexa fluor (488, 555, and 647) conjugated donkey anti-rabbit, and goat anti-mouse (Invitrogen).

### Immunostaining of co-cultures

Co-cultures were washed with 1xPBS for 5 minutes and fixed with 4% paraformaldehyde (PFA) for 12-15 minutes. PFA was then removed, and cells were washed with 1xPBS 2 times for 5 minutes. Then slides were blocked for 30 minutes with 1% BSA in PBT (PBT - PBS with 0.1% Triton). Then enough primary antibody was prepared and diluted in 0.1% BSA in PBT. The blocking solution was removed, and the primary antibody solution was added and incubated overnight at 4 degrees celsius in a humidified chamber. The next day, the primary solution was removed, and slides were washed 3 times for 5 minutes with PBT at room temperature. Secondary antibodies were prepared in PBT and were filtered through a syringe filter (Millipore Millex-GV 13mm x 0.22um). The secondary antibody solution was added to the slides and incubated for 60 minutes at room temperature in a humidified chamber. After 1 hour, slides were washed 3 times for 5 minutes with PBT at room temperature. Slides were then mounted with Vectashield and sealed with nail polish **(Sup.** Figure 5B).

Rabbit anti-synaptophysin (Thermo Fisher; 1:500 dilution) was used as the primary antibody for motor neurons in co-cultures. Alpha-bungarotoxin 555-conjugate (Thermo Fisher; 1:500 dilution) was added with the secondary antibody solution.

### Analysis of Immunostaining

All immunostaining images were processed with Fiji (ImageJ). Synapses were blindly determined with the overlap of *α*BTX with SYP. The number of synapses in different conditions was statistically analyzed with an unpaired t-test assuming both populations had the same standard deviations **(Sup.** Figure 5B). For all statistical tests, a value of *p* ≤ 0.05 was considered statistically significant, with *p* ≤ 0.05 denoted as *, *p* ≤ 0.01 denoted as **, *p* ≤ 0.001 denoted as ***, *p* ≤ 0.0001 denoted as ****, and ns is not significant.

## DECLARATION

### ETHICS APPROVAL AND CONSENT TO PARTICIPATE

All experiments were carried out according to human stem cell (hESCRO) protocols that were approved by Case Western Reserve University.

## CONSENT FOR PUBLICATION

Not applicable.

## ACKNOWLEDGEMENTS

The authors would like to thank Dr. Lorenzo Puri from the Sanford Burnham Prebys Medical Discovery Institute for the gift of the MyoD and BAF60C constructs and Dr. Fred Gage from the Salk Institute for the gift of the SYN::hChR2:YFP construct. The HB9 and ISL1 antibodies developed by the Jessell lab and the Myogenin antibody developed by the Wright lab were obtained from the Developmental Studies Hybridoma Bank, created by the NICHD of the NIH and maintained at The University of Iowa, Department of Biology, Iowa City, IA 52242

## AVAILABILITY OF DATA AND MATERIALS

All material and data generated here will be available upon request.

## AUTHORS’ INFORMATION

**Daniel Chen and Bianca de Freitas Brenha**, Department of Genetics and Genome Sciences, Case Western Reserve University, Cleveland, OH USA. **Polyxeni Philippidou**, Department of Neurosciences, Case Western Reserve University, Cleveland, OH USA **Ashleigh E. Schaffer**, Department of Genetics and Genome Sciences, Case Western Reserve University, Cleveland, OH USA, Center for RNA Science and Therapeutics, Case Western Reserve University, Cleveland, OH USA and **Helen C. Miranda** Department of Genetics and Genome Sciences, Case Western Reserve University, Cleveland, OH USA, Department of Neurosciences, Case Western Reserve University, Cleveland, OH USA Center for RNA Science and Therapeutics, Case Western Reserve University, Cleveland, OH USA

## AUTHORS’ CONTRIBUTIONS

HM and DC designed experiments. HM, DC, and AS prepared the manuscript. DC generated PSC-derived skeletal muscles and motor neurons. DC developed PSC-derived NMJs. PP performed PSC-derived NMJ immunostaining. BB performed microscopy for blinded counting of NMJs. DC performed and analyzed the functional electrophysiological recording of PSC-derived NMJs. All authors read and approved the final manuscript.

